# Humidity-dependent structural adaptations of *Drosophila melanogaster* hygrosensilla

**DOI:** 10.1101/2024.11.19.624428

**Authors:** Ganesh Giri, Anders Enjin

## Abstract

Understanding how organisms detect environmental humidity remains a fundamental problem in sensory biology. While specialized sensory neurons in insect antennae can detect changes in humidity, the mechanism underlying this ability is not fully understood. Here, we present an integrated approach combining precise humidity control, rapid cryo-preservation, and serial block-face scanning electron microscopy (SBF-SEM) to investigate the ultrastructure of hygrosensilla in the vinegar fly *Drosophila melanogaster*. We developed a deep learning-based segmentation pipeline to analyze three-dimensional structural features of sensilla exposed to different humidity conditions at stable temperature. Our analysis reveals consistent differences in sensilla width between high (80% RH) and low (26% RH) humidity conditions across all chambers of the sacculus. Additionally, we identified chamber-specific patterns in sensilla tapering, indicating specialized structural adaptations across different sensilla populations. The observed structural changes suggest a potential role for mechanical transduction in humidity sensing. This study establishes a technical framework for high-resolution analysis of sensory organs while providing new insights into the structural basis of humidity detection. Our findings advance our understanding of how specialized sensory organs might transduce environmental signals into neural responses.

## Introduction

Humidity is a fundamental environmental factor that influences the ecology and behaviour of terrestrial organisms [1–3]. The ability to sense and respond to changes in ambient moisture, known as hygrosensation, is crucial for a wide range of species. Small ectotherms, such as insects, are particularly sensitive to humidity due to their high surface area to volume ratio, which makes them vulnerable to water loss [4,5]. Numerous species have evolved to utilize local variations in humidity as vital cues for essential behaviours such as foraging, reproduction, and habitat selection [6–13]. Despite its importance in the biology of diverse organisms, our understanding of the mechanisms underlying hygrosensory transduction remains incomplete, presenting a significant gap in our knowledge of sensory biology.

In insects, humidity is sensed via specialized sensory hairs called hygrosensilla, typically found on the antennae [14,15]. These hygrosensilla are often located in protected areas, such as invaginations or grooves, shielding them from wind and mechanical disturbance, presumably to provide a more accurate reading of environmental humidity. For example, in the vinegar fly *Drosophila melanogaster*, the hygrosensilla are located in a relatively large invagination on the posterior side of the antenna called the sacculus [16–18]. The typical hygrosensillum houses a triad of sensory humidity receptor neurons (HRNs): a moist neuron that increases its firing rate in response to rising humidity, a dry neuron with an inverted response profile, and a hygrocool neuron sensitive to cooling [19,20].

The unique features of hygrosensilla present a challenge to understanding hygrosensory transduction. These features include their pore-less structure, protected location, and composition of both hygrosensory and thermosensory neurons. Furthermore, the ambiguous nature of water vapor as a ‘ligand’ for a receptor submerged in a water-based solution (lymph) complicates this understanding. Three main hypotheses have been proposed to explain the mechanism of hygrosensation: mechanosensation, osmosensation, or thermosensation [21,22]. The mechanosensation model suggests that HRNs detect structural changes in hygrosensilla through mechanically-gated channels. The osmosensation hypothesis proposes that HRNs detect changes in ion concentration in the sensillum lymph due to water evaporation. The thermosensation model suggests that HRNs detect the rate of cooling caused by evaporative cooling. Electrophysiological studies have revealed important insights into how the HRNs might work together in to detect changes in humidity. Recordings from the moist and dry cells show a double dependency on both temperature and humidity that cannot be reduced to a common parameter such as evaporative cooling [22]. For example, in cockroach hygroreceptors, when water vapor pressure is oscillated at different temperature levels, the discharge rate of moist cells increases with rising vapor pressure while dry cells show the opposite pattern [22]. Importantly, the higher the temperature of these oscillating changes in vapor pressure, the stronger the oscillating responses of both moist and dry cells. The fact that both cells’ responses are modulated by humidity and temperature suggests that evaporative cooling alone cannot fully explain hygrosensation. Moreover, when the water vapor content of the air remains constant, but temperature rises, relative humidity decreases, yet impulse frequency in both moist and dry cells increase with rising temperature [22]. This positive temperature coefficient of the hygroreceptors’ responses to changes in relative humidity contradicts a purely mechanical transduction model. Together, these findings indicate that hygroreceptors likely use multiple mechanisms, combining temperature, osmotic and/or mechanical inputs in ways that cannot be explained by any single transduction model. Furthermore, studies in *C. elegans* have shown that hygrosensation requires a combination of distinct mechanosensitive and thermosensitive pathways, suggesting this dual-input strategy may be evolutionarily conserved across species that lack specialized hygroreceptors [23]. Similarly, in human wetness perception, both mechanical pressure and cold sensation through TRPM8 channels is required [24].

Several studies have suggested that mechanosensation is key feature of hygrosensation also in insects, however demonstrating this morphologically has proved difficult [25–27]. Studies on the domestic silk moth *Bombyx mori* found humidity-induced morphological changes, but only after extended periods of dry and moist adaptation, preventing insight into immediate structural responses [28]. A different approach using in-situ atomic force microscopy on honey bee *Apis mellifera* hygrosensilla revealed only limited lateral structural changes, possibly due to technical constraints in detecting fine-scale alterations [29]. Here, we use rapid plunge-freezing combined with serial block-face scanning electron microscopy (SBF-SEM) to investigate rapid structural responses of *D. melanogaster* hygrosensilla to humidity change. This approach allows us to capture and analyze three-dimensional structural changes across the entire sensillum, revealing consistent differences in sensilla dimensions between high and low humidity conditions and providing new insights into how these specialized sensory organs might transduce environmental signals into neural responses.

## Materials and methods

### Sample Preparation

Female *w^1118^* flies, aged 7-14 days and with intact appendages and antennae, were selected for the experiment. To preserve the flies under a specific humidity condition, Flies were attached to forceps positioned within the humidity-controlled chamber of a Vitrobot Mark IV system (FEI, Denmark) and were held at the desired humidity level for approximately 45 seconds before being plunged into liquid ethane. The setup included a crucible on a platform beneath the chamber, with liquid nitrogen in the outer cavity and liquid ethane in the inner cavity. This rapid freezing process ensured that the fly was preserved under the designated humidity levels (Figure 1A). Two humidity conditions were used: high humidity at 80% relative humidity (RH) and low humidity at 26% RH both maintained at a room temperature of 22° C. Following plunge freezing, samples were transferred to a freeze substitution medium containing 0.1% tannic acid and 0.5% glutaraldehyde in acetone and incubated at −90°C for 96 hours. Samples were then washed four times with anhydrous acetone to remove the initial freeze substitution medium. The acetone wash was replaced with a fresh solution of 2% osmium tetroxide (OsO₄) in anhydrous acetone. The freeze substitution process was continued with this OsO₄ solution for an additional 28 hours. Osmium tetroxide was used to stain the samples, providing contrast for electron microscopy by binding to lipids and preserving cellular structures. After staining, the temperature was gradually raised over 14 hours to −30°C, where the samples were held for 16 hours. To remove the OsO₄ and any remaining residues, the samples were washed four more times with anhydrous acetone. The temperature was then further increased over 4 hours to 1°C. Dehydration was performed through a graded ethanol and acetone series, with each step lasting 10 minutes. The sequence included 70% ethanol on ice, 90% ethanol on ice, two changes of 100% ethanol on ice, followed by 100% acetone with ice-cold acetone at room temperature, and finally, 100% acetone at room temperature. After dehydration, the samples were embedded in Epon for further processing.

**Figure 1:**
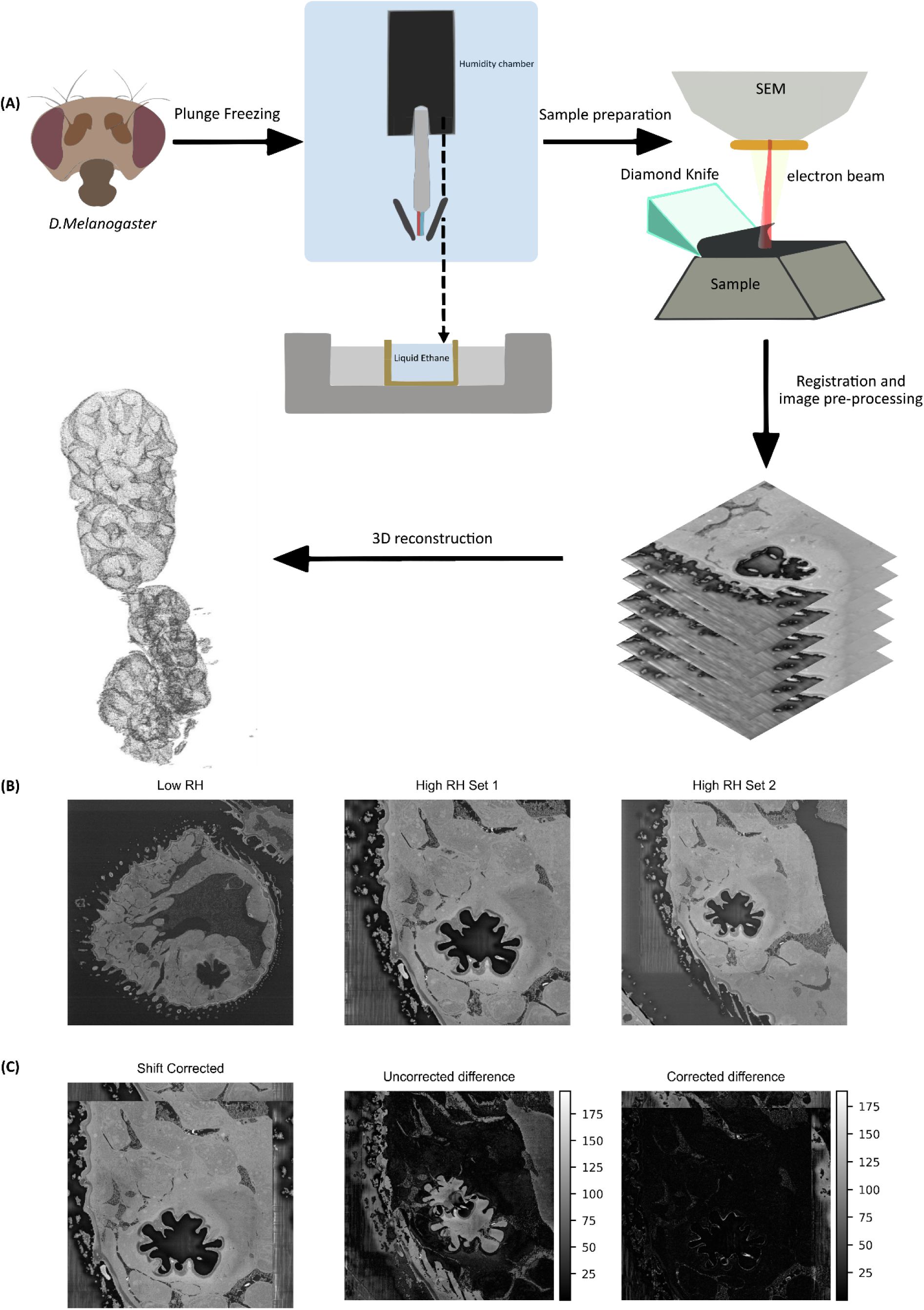
Workflow for sample preparation, imaging, and segmentation of *D. melanogaster* sacculus under varying humidity conditions. (A) Schematic of the plunge-freezing protocol: flies were placed in a humidity-controlled chamber (26% or 80% RH) for 30 seconds, then plunge-frozen in liquid ethane to preserve their structural state. The sample was stained, embedded in epoxy resin, and trimmed to expose the antenna, which was imaged in its entirety using SBF-SEM. Segmentation of the sacculus from these images was achieved using a U-Net model, enabling 3D reconstruction. (B) Sample images acquired at low relative humidity (Left) and high relative humidity (Center and Right). (C) High RH sample images (Set 2) adjusted to match Set 1 resolution, with difference images showing Set 1 vs. Set 2 before (Center) and after (Right) shift correction and resolution correction.

### Imaging

The embedded resin block was positioned and trimmed to achieve an optimal angle for acquiring antennal sections. SBF-SEM was performed on the trimmed samples using the Teneo VolumeScope (FEI, Denmark). For the sample prepared at 80% RH (high RH dataset), a total field of view of 42.5 µm × 42.5 µm, with a total depth of 14.46 µm was achieved with the imaging resolution of 10.3760 nm × 10.3760 nm in the X/Y plane and 30 nm in the Z-axis (High RH Set 1), with the sample sliced into 482 sections of 30 nm thickness each. Beam parameters included an energy of 2.28 keV, an electron dose of 69.57 e/nm², a beam current of 400 pA, and a dwell time of 3 µs. Imaging was conducted under low vacuum conditions, using a VsGAD detector with line and frame integration set to 1. The tile resolution was 4,096 × 4,096 pixels. Multi-Energy Deconvolution (MED) was applied to enhance contrast by combining images captured at different energy levels, effectively reducing noise and improving structural detail. Following an interruption in the imaging process, the sample was repositioned, and imaging was resumed. This led to a change in imaging parameters. The field of view was expanded to 60.0 µm × 60.0 µm, covering a total depth of 30.03 µm with a resolution of 14.6484 nm × 14.6484 nm in the XY plane (High RH Set 2). The resolution along the Z-axis remained at 30 nm. A total of 1,001 slices were acquired under similar beam conditions, with an electron dose of 34.91 e/nm². The same detector and imaging settings were used as before.

For the sample prepared at 26% RH (low RH dataset), the imaging condition was similar without any interruptions during image acquisition. A field of view of 111.0 µm × 111.0 µm and a depth of 104.16 µm was achieved with a resolution of 10.839 x 10.839 nm in the XY plane and 30 nm in the Z plane. A total of 3472 slices were collected with an energy of 2.27KeV and an electron dose of 10.62 e/nm² with a dwell time of 1 µs.

### Image rescaling and alignment for high RH dataset

To align and analyse the images from the high-humidity dataset (80% RH), it was necessary to account for the differences in resolution between two sets of images obtained during the experiment (High RH Set 1 and High RH Set 2). The first set of images had a pixel size of 10.3760 nm × 10.3760 nm in the X/Y plane, while the second set had a larger pixel size of 14.6484 nm × 14.6484 nm. To account for this discrepancy, scale factors were calculated based on the pixel size ratio between the two sets. The images from the second set were rescaled according to the scaling factor to get a resolution of 10.3760 nm × 10.3760 nm in the X/Y plane. Rescaling was done with the help of rescale function from skimage library, with anti-aliasing applied to minimize artifacts [30]. Following rescaling, the images from the second set were cropped or padded to match the exact dimensions of the first set

For alignment, the last image from High RH Set 1 and the first rescaled image from High RH Set 2 were compared using phase cross-correlation, implemented via phase_cross_correlation function from the skimage library. This method estimated the translational shift between the two images, which was then applied to the entire second set using a Fourier shift-based method, ensuring subpixel accuracy. After applying the shift, the images were normalized to be in the range of 0-255 to maintain consistent intensity levels throughout the dataset. The final aligned and normalized images were saved for further analysis.

### Generating image labels

A custom Python script was used to randomly select 337 images from both the High and Low RH datasets, ensuring an equal proportion of images from each condition. The sacculus outlines were manually traced using the drawing tools in IMOD software [31]. These traced outlines were then converted into a binary label stack using the imodmop function to generate binary labels, with the mask value set to 1.

### Data augmentation

To increase the diversity of training data for the machine learning model, data augmentation was performed on the labelled images. For each labelled image, four new augmented versions were generated: vertically flipped, horizontally flipped, and rotated by 45 and 130 degrees. The augmented images and their corresponding labels were saved in separate directories for further use in training the model.

### Segmentation

A deep learning-based approach was implemented using a U-Net architecture to segment the sacculus from the acquired images [32]. The images and their corresponding labels were resized to a dimension of 256×256 pixels and then normalized to have pixel values in between 0 and 1. The pre-processed data was then split into training, test and validation sets in the ratio of 90:5:5.

The U-Net model was compiled using Adam optimizer with binary cross entropy as the loss function. Accuracy and Intersection over union (IoU) were used as metrices to evaluate the training of the model. Since the region of interest corresponding to the sacculus was much smaller in area compared to the background, IoU metric was given more importance compared to the accuracy metric during the evaluation of the model.

Two callback functions from the tensor flow library were implemented to enhance the performance of the trained model [33]. The first was ModelCheckpoint, which saved the model with the best IoU score during the training to ensure that the model with the highest overlap between the predicted mask and ground truth was saved. The other callback being ReduceLROnPlateau which dynamically reduced the learning rate whenever the validation loss plateaued, aiding in more effective convergence of the model.

Using the trained model, predicted labels segmenting the sacculus were generated for both the dry and high RH dataset. The predicted labels had a size of 256 x 256 pixels.

### Generating 3D Structure of the sacculus

Contours were extracted from the predicted labels using contour detection function from the OpenCV library with RETR_TREE as the retrieval mode [34]. This allowed for the detection of all the nested contours within the predicted labels. Detected contours were stacked and assigned a Z coordinate according to its position in the image sequence. The contour stack was then rescaled to match the original image dimension and to account for the physical resolution of the data set. The scaled contour points were then used to generate a point cloud which visualized the 3-dimensional structure of the segmented sacculus.

### Isolation and Alignment of the sensilla

The point cloud was visualized using the Open3D library, which facilitated the identification and localization of individual sensilla [35]. The identified sensilla were then manually cropped using the draw_geometries_with_editing function from Open3D.

The cropped sensilla varied in orientation depending on their position within the sacculus, which made measuring their dimensions challenging. Therefore, orienting them along a specific axis was important. To standardize orientation, we selected one representative sensillum from each chamber (Chambers I and III) and centered its point cloud at the origin by subtracting the centroid coordinates from each point. Principal Component Analysis (PCA) was then applied to this centered point cloud to identify its principal axes. In sensilla with a large height-to-width ratio, such as those in chambers I and III, the axis with the greatest variance corresponded to the sensilla’s length. This axis was designated as the principal axis. The angle between this principal axis and the Z axis was then calculated, and a rotation was applied to orient the sensillum along the Z axis. This oriented sensillum served as the target for aligning the remaining sensilla within the chamber. A RANSAC algorithm [36] was implemented to perform global registration, aligning each remaining sensilla’s point cloud with that of the target. First, the point clouds were downsampled, and then a global registration process was used to compute the optimal transformation matrix, achieving the best alignment with the target

However, in the case of Chamber II, the PCA analysis did not yield the correct orientation, as the chamber’s length and width were similar, causing the axis of maximum variance not to correspond with the length of the sensilla. To address this, a conical point cloud matching the dimensions of the sensilla in Chamber II was created and oriented along the Z-axis. This synthetic point cloud served as the target for alignment. Using a RANSAC-based approach, the sensilla in Chamber II were aligned to the target point cloud, ensuring accurate and consistent orientation for all sensilla within the chamber.

### Width measurement

The aligned point clouds were converted into a 3D mesh using a Delaunay triangulation approach. The meshes were then sliced at different heights to generate 2D cross sectional slices. For each slice, the width of the sensilla was determined by calculating the maximum distance between the extreme points along the XY plane. This effectively captured the widest span of the structure at that specific height. This process was repeated across multiple slices, providing a width profile along the height of the sensilla.

Noise in width measurement was reduced by applying a weighted moving average (WMA) filter to the width profile. After smoothing the width of the corresponding height values were plotted to analyse the sensilla’s width distribution along its length.

### Data Analysis

A mixed-effects model from statsmodels was implemented to examine the relationship between sensilla width and height, while accounting for differences between the high and low RH dataset [37]. The model included a fixed effect for the interaction between height and group (humid vs. dry) to capture the influence of environmental conditions on width. Additionally, a random effect for individual sensilla was incorporated to account for inherent variability between sensilla. This random effect allowed the model to adjust for baseline differences in width across individual sensilla, ensuring that the analysis focused on the effects of the humidity conditions rather than on the natural variation between sensilla

After fitting the model, predictions were generated for the width of sensilla across a range of heights for both groups. The model results were used to calculate 95% confidence intervals around the predicted widths to evaluate the uncertainty in the estimates.

The dimensions of the sensilla were further compared using its full width half maximum (FWHM). FWHM for each sensilla was determined by identifying the height of individual sensilla and then locating the corresponding width at half the value of the determined height, thereby providing a standardized method to compare the overall size of the sensilla. To assess differences between groups, the FWHM distributions were compared using a non-parametric Mann-Whitney U test.

## Results

### Structural characterization of sensilla across humidity conditions

Using SBF-SEM, we investigated the ultrastructural organization of hygrosensilla in the antennal sacculus of *D. melanogaster* under two distinct humidity conditions. Flies were exposed to either high (80% RH) or low (26% RH) humidity at 22°C for approximately 45 seconds and then rapidly plunge-frozen in liquid ethane (Figure 1A). This approach allowed us to capture and preserve the sensilla in their native state under a specific humidity condition. The SBF-SEM imaging provided comprehensive volumetric data of the sacculus, covering dimensions of 42.5 × 42.5 × 14.46 µm (High RH Set 1) and 60.0 × 60.0 × 30.03 µm (High RH Set 2) at resolutions of 10.38 and 14.65 nm/pixel respectively. For the low humidity condition, we acquired a larger volume of 111.0 × 111.0 × 104.16 µm at 10.84 nm/pixel resolution (Figure 1B-C). These high-resolution datasets enabled three-dimensional reconstruction of the complete sacculus structure, including all three chambers and their associated sensilla.

### Segmentation of sensilla

To systematically analyse the structural properties of individual sensilla, we developed a deep learning-based U-Net segmentation model to accurately identify and isolate sensilla from the sacculus. The model achieved high classification accuracy, with a 95% overall accuracy throughout the training phase and an Intersection over Union (IoU) metric of 0.854 (Figure 2A), a robust measure of segmentation performance, particularly in regions with a small region of interest relative to the background. Despite the complex structure of the sacculus and the small size of the sensilla, the model performed effectively, allowing for reliable identification of the sacculus across both the Heigh and Low RH datasets (Figure 2B).

**Figure 2:**
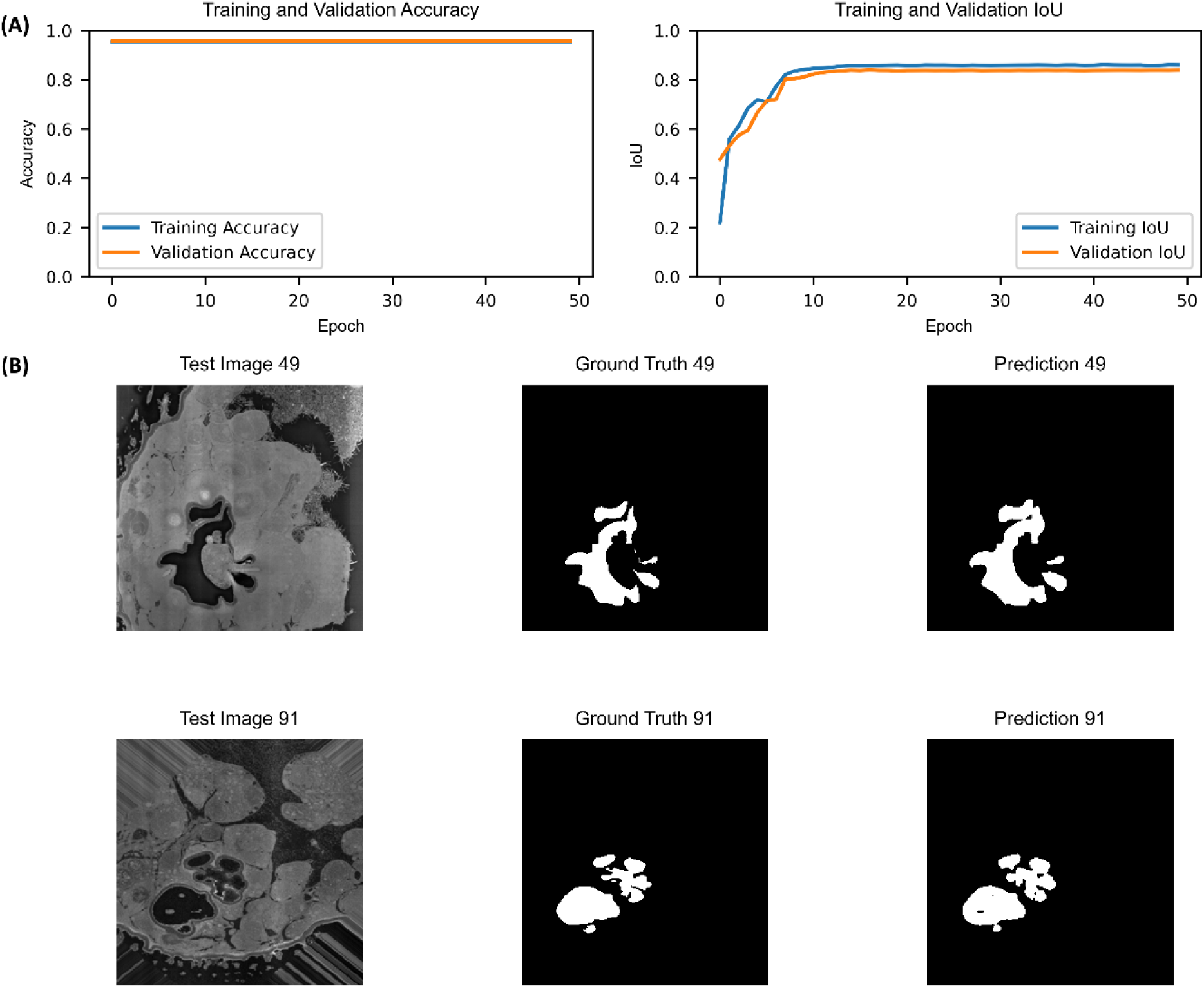
Evaluation and application of the U-Net model for sacculus segmentation. (A) Model performance metrics, showing Accuracy (Left) and Intersection over Union (IoU), for both training and validation datasets. (B) Segmentation results using the trained U-Net model on test data, displaying the test input (Left), the corresponding ground truth (Center), and the model’s prediction (Right).

With the contours generated by using the segmented image, we recreated the 3D structure of the sacculus (Figure 3 A-B) and identified four distinct types of sensilla distributed across the three chambers of the sacculus. In Chamber 1, the sensilla appeared elongated and slender, maintaining a consistent taper toward the tip. In contrast, sensilla in Chamber 2 were characterized by a shorter and wider structure, with a broader base and less pronounced tapering. Chamber 3 featured two types of sensilla, both elongated; however, the sensilla in the lower section of this chamber were notably wider than those found in the upper section (Figure 4A-C). These structural variations were present across both humidity conditions but showed noticeable differences in dimensions when compared between the High and Low RH samples.

**Figure 3:**
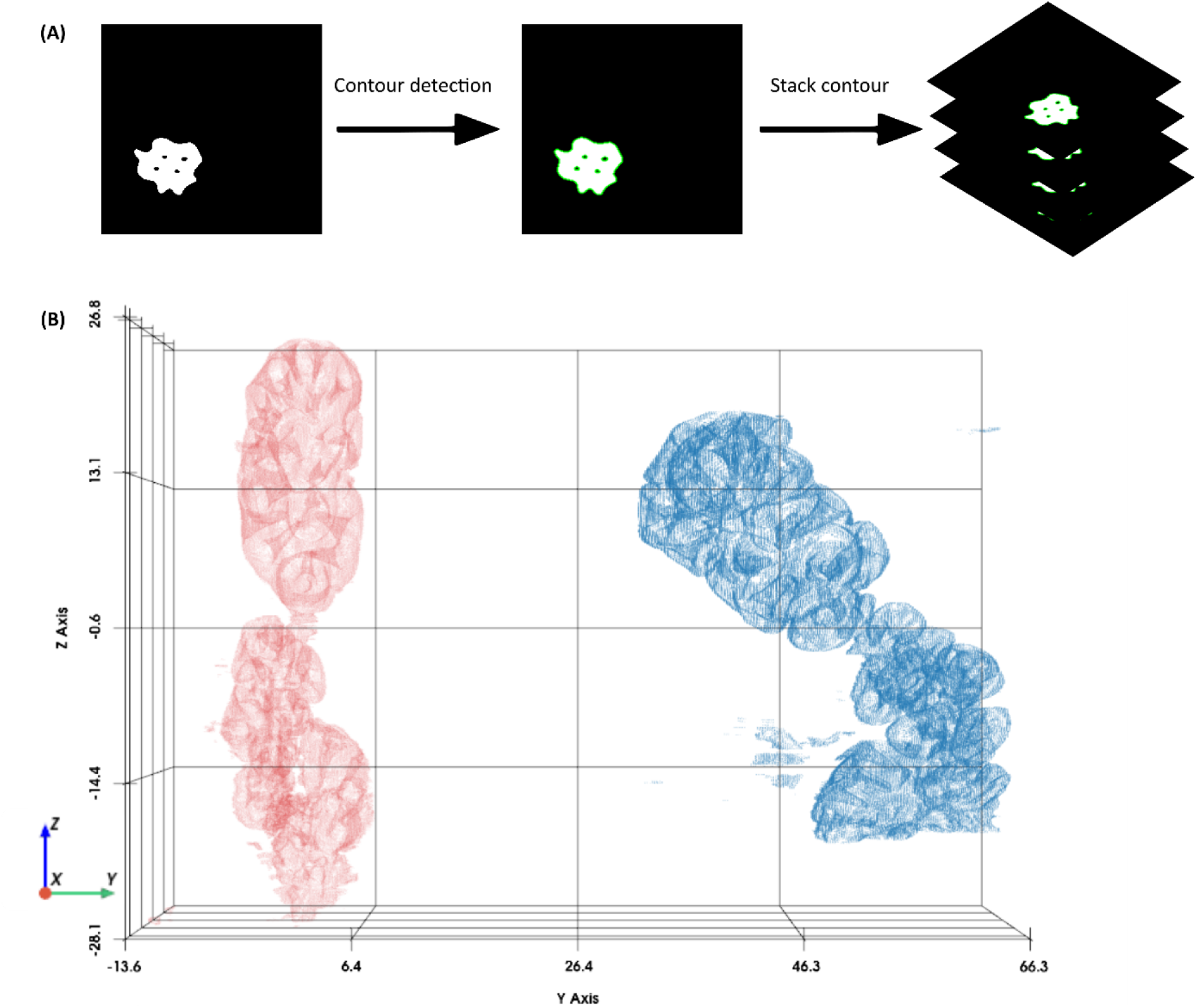
Sacculus structure representation through contour detection and point cloud visualization. (A) Contour detection was performed on each segmented image in the stack. (B) The detected contours were converted into point clouds to visualize the sacculus structure. The red point cloud represents the sacculus from the low humidity (RH) sample, and the blue point cloud represents the sacculus from the high humidity (RH) sample.

**Figure 4:**
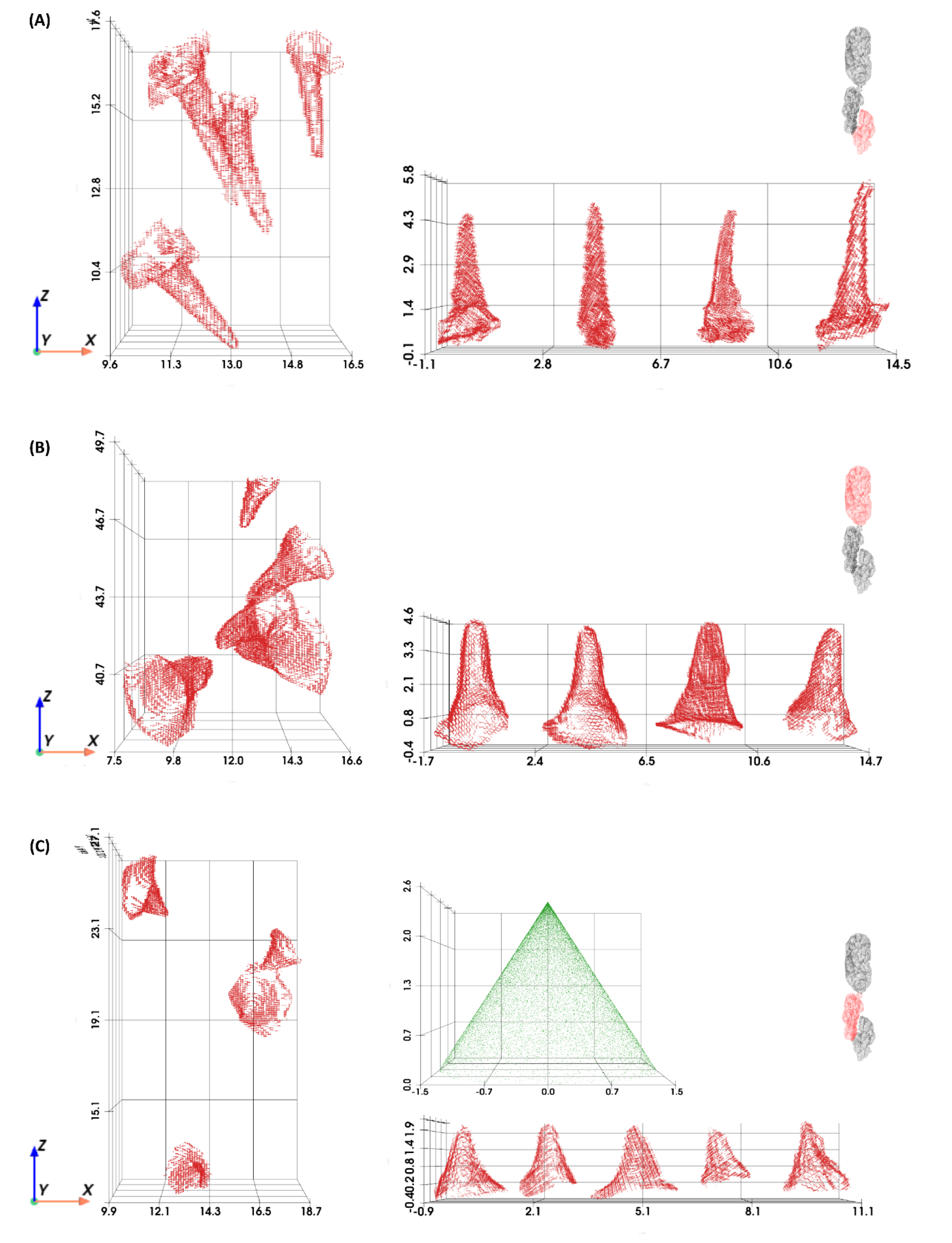
Isolated sensilla from each chamber, shown in their original orientation within the sacculus (Left) and after alignment for dimensional measurement (Right). (A) Chamber 1, (B) Chamber 3, and (C) Chamber 2. For Chamber 2, the green conical point cloud was used as a reference to align sensilla accurately for measurement

In the High RH sample, our segmentation successfully identified five distinct sensilla in both Chambers 1 and 2, while in the Low RH sample, four sensilla were clearly identified in each of these chambers. Additional sensilla were partially visible but could not be fully reconstructed due to incomplete segmentation. Chamber 3 provided the most complete reconstructions in both the samples, with multiple sensilla

### Quantitative Analysis of Sensilla Width and Height

Quantitative analysis of sensilla dimensions was carried out following image segmentation and realignment. To ensure accuracy in our measurements, we utilized image scaling techniques to account for differences in resolution between the image sets acquired before and after the imaging interruption in the high humidity condition. This allowed us to obtain consistent measurements across all samples, enabling a robust comparison of sensilla dimensions under varying humidity conditions.

In the high humidity dataset (80% RH), the average width of the sensilla in Chamber 1 was 3.36 µm with a standard deviation of 0.13 µm. This width was 0.77 µm greater than the mean sensilla width in the Low RH dataset (26% RH), where the average width measured 2.59 µm. A similar pattern was observed in Chambers 2 and 3, where sensilla in the high humidity samples consistently exhibited wider dimensions than their counterparts in the low humidity condition (Figure 5A-B).

**Figure 5:**
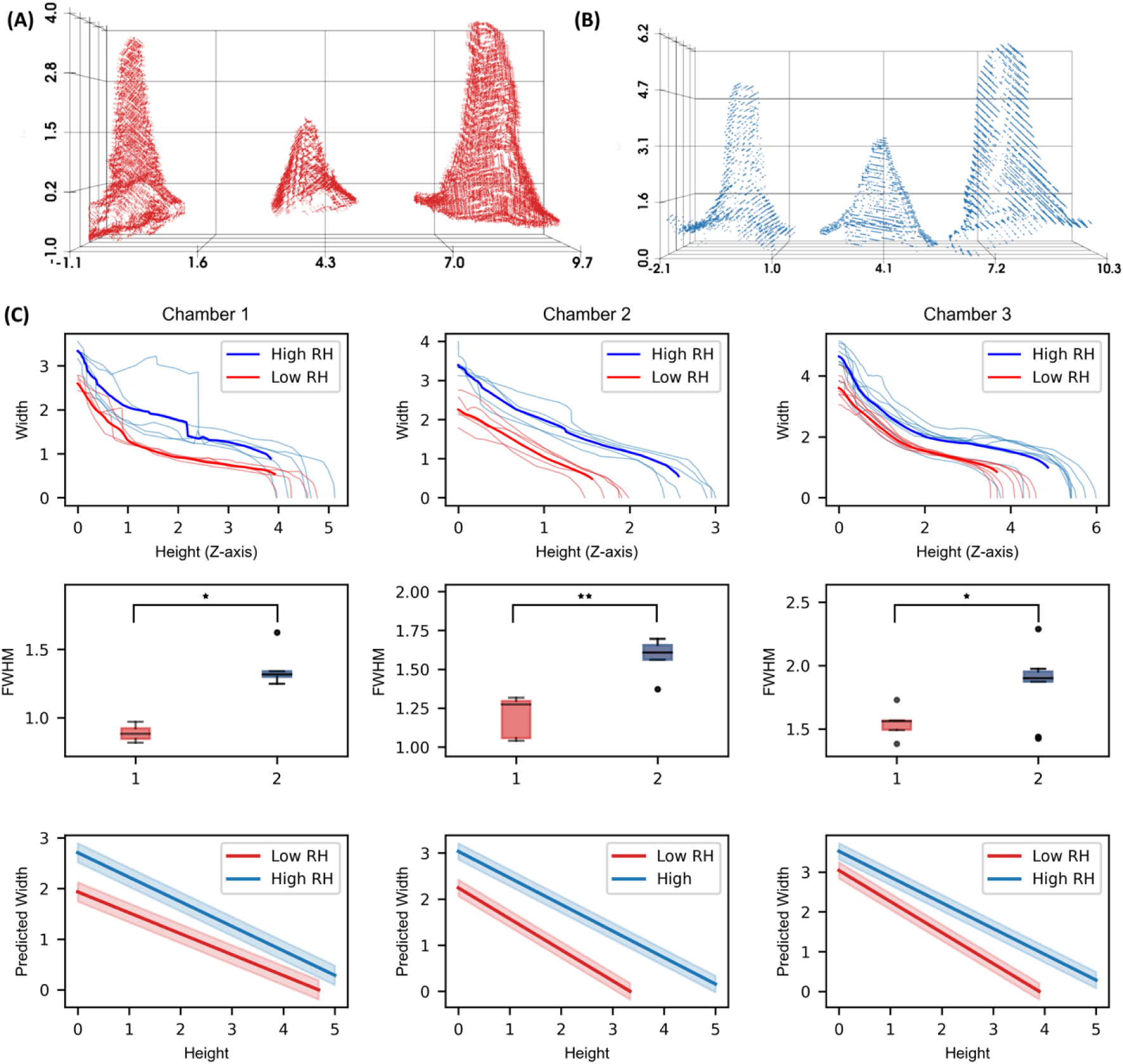
Structural comparison of *D. melanogaster* sensilla across humidity conditions. (A) Sensilla point clouds from low humidity (26% RH) and (B) high humidity (80% RH) samples, oriented and aligned for Chambers 1, 2, and 3 from left to right. (C) Top: Length vs. width profiles for each identified sensillum in Chamber 1 (Left), Chamber 2 (Center), and Chamber 3 (Right), with low humidity samples in red and high humidity samples in blue; darker curves represent the mean length vs. width profiles for each condition. Middle: Full width at half maximum (FWHM) values for each identified sensillum, calculated by determining the height of each sensillum and locating the corresponding width at half this height; statistically significant differences (Mann Whitney U test, * p < 0.05, ** p <0.01, *** p< 0.001) are observed between high and low RH conditions across all chambers. Bottom: Mixed-effects model of the length vs. width profiles. Solid lines represent calculated profiles, with shaded areas indicating the 95% confidence intervals. Notably, in Chamber 1, high humidity sensilla display a steeper taper (slope: −0.472) compared to low humidity (slope: −0.403). In contrast, sensilla in Chambers 2 and 3 exhibit a steeper taper under low humidity (slopes: −1.120 and −0.652, respectively) than under high humidity (slopes: −0.959 and −0.541).

The most pronounced difference in the magnitude of median full-width at half maximum (FWHM) was observed in Chamber I, with a 0.4 µm increase in sensilla width in the high RH dataset compared to the low RH dataset (p< 0.05). In Chambers 2 and 3, the magnitude of this difference was slightly less pronounced but remained statistically significant, with the high humidity sensilla showing an increase of 0.33 µm in both the chambers (p< 0.01 for Chamber II and p<0.05 for Chamber III) (Figure 5C). This indicates a similar pattern of expansion in response to increased environmental moisture in both Chamber II and III.

### Height-Width Relationship Across Humidity Conditions

To further explore the relationship between sensilla width and height under different humidity conditions, we employed a mixed-effects model, which allowed us to account for the inherent variability between individual sensilla while analysing the effects of humidity. The model revealed significant differences in the rate of decline in sensilla width with increasing height between the high and low humidity conditions.

In Chamber 1, sensilla in the high humidity condition exhibited a steeper decline in width with increasing height, with a slope of −0.472 compared to −0.403 in the low humidity condition. This suggests that sensilla under high humidity were not only wider at the base but also showed a sharper tapering toward the tip. This structural adaptation may play a role in optimizing sensitivity of the hygrosensory neurons housed within the sensilla.

Conversely, in Chambers 2 and 3, the decline in sensilla width with height was more pronounced in the low humidity condition. In Chamber 2, the slope was −1.120 under dry conditions compared to −0.959 in the high humidity samples, indicating a steeper reduction in width towards the tip of the sensilla in the low humidity environment. Similarly, in Chamber 3, the slope for the dry condition was −0.652, compared to −0.541 for the humid condition (Figure 5B). These findings suggest that while sensilla in humid conditions are generally wider, they maintain a more gradual taper, potentially facilitating a more nuanced detection of humidity changes across varying sensilla types.

## Discussion

### Structural plasticity of hygrosensilla

Here, we present a comprehensive ultrastructural analysis of humidity-dependent changes in *D. melanogaster* antennal hygrosensilla using SBF-SEM under precisely controlled humidity and temperature conditions. This approach enables high-resolution, three-dimensional reconstruction of sensilla structure while maintaining their native state under specific humidity conditions. Our results reveal that hygrosensilla exhibit structural plasticity in response to ambient humidity, with sensilla in high humidity conditions (80% RH) showing consistently greater widths compared to those in low humidity conditions (26% RH). This dimensional difference was most pronounced in Chamber 1, where sensilla demonstrated a 0.77 µm larger width under high humidity, while sensilla in Chambers 2 and 3 showed similar but less dramatic changes.

The chamber-specific structural profiles suggest distinct adaptations across different sensilla populations. Under high humidity, the sensilla in Chamber 1 displayed a sharper taper from base to tip, which likely concentrates the maximum mechanical pressure changes toward the base. In contrast, sensilla in Chambers 2 and 3 exhibited more pronounced tapering toward the tip under low humidity, possibly optimizing sensory function by focusing pressure changes toward the tip in drier conditions. These non-uniform, localized hygroscopic changes in the sensilla suggest that the cuticular wall’s water-absorbing properties contribute to chamber-specific swelling and shrinking patterns.

The analysis of these structural adaptations was facilitated by our U-Net-based segmentation approach, which achieved an IoU score of 0.854 in detecting the sacculus within the antennal structure. However, as sensilla dimensions were measured from separate individuals-one prepared at high humidity and one prepared at low humidity conditions, the influence of individual variability cannot be fully excluded. While the structural differences observed are consistent across the two samples, further studies involving larger sample size is needed to confirm that these morphological differences are driven solely by humidity rather than being individualistic variations. In addition, our current methodology, while providing detailed structural information, represents static snapshots of what is likely a dynamic mechanical response. Nevertheless, the consistent patterns observed across multiple sensilla within each specimen support a model where structural adaptation plays a role in mediating *D.melanogaster’s* humidity sensitivity.

### Mechanistic Basis of Humidity Detection

Understanding the mechanism of hygrosensory transduction presents unique challenges due to the specialized anatomy of hygrosensilla. Unlike olfactory sensilla that detect airborne molecules through pores in their cuticle, hygrosensilla have a poreless structure and are therefore likely to detect a physical stimulus rather than a chemical [14]. Our ultrastructural analysis provides support for mechanosensation as a component. The increase in sensilla width observed under high humidity conditions suggests that the sensilla undergo structural changes in response to environmental humidity. Since HRNs protrude their sensory cilia into the hygrosensillum and maintain close contact with the cuticular wall [16], these dimensional changes could activate mechanosensors in the plasma membrane of the HRNs. Transcriptomic analysis of HRNs has revealed distinct molecular signatures that underscore the intimate relationship between sensory neurons and their associated sensilla structures [38]. Dry- and moist-sensing HRNs exhibit differential expression patterns of cuticule-associated proteins potentially forming a link between the membrane of the HRN and the cuticle, vermiform expressed in dry neurons and fred is expressed in moist neurons [39–41]. Both proteins are integral to chitin metabolism and interaction, suggesting that specialized cuticular architecture may be fundamental to hygrosensory function. Furthermore, these neuronal populations show complementary expression of nicotinic acetylcholine receptor subunits α6 and α7. These receptors are particularly significant as their orthologs have been implicated in mechanotransduction [42,43], potentially indicating a mechanosensory component in hygrosensation.

If humidity-induced expansion and contraction of the sensillum cuticle drives neuronal activation, one would expect the responses to track relative humidity (RH), as any hygroscopic mechanical changes, such as that of a human hair, would correlate with RH [29]. However, electrophysiological studies show that both moist and dry cells display a positive temperature coefficient, their responses to humidity fluctuations increase with rising temperature, contrary to what would be expected from simple RH sensors [21]. This apparent contradiction can be resolved by considering the role of ephaptic coupling between the compartmentalized hygrosensory neurons. Recent work has demonstrated that sensory neurons housed in the same sensillum can inhibit each other through direct electrical field effects mechanisms, without requiring synaptic connections [44–46]. We hypothesize that the hygrocool cell, consistently present alongside the dry and moist cells across insect species, may use ephaptic inhibition to modulate the responses of its neighbouring mechanosensitive neurons. As temperature decreases, the hygrocool cell’s activity increases, leading to stronger ephaptic inhibition of both the dry and moist cells [47]. This explains why their humidity responses are reduced at lower temperatures, despite the mechanical response of the sensillum to RH remaining constant. While speculative, this model accounts for a functional explanation for the highly conserved grouping in hygrosensilla of these three cell types, the hygrocool cell acts as a temperature-dependent gain control mechanism for the mechanosensitive dry and moist cells through ephaptic coupling.

### Temporal Dynamics and Spatial Organization of Humidity Sensing

The structural adaptations we observe in hygrosensilla must support rapid sensory responses, as flies exhibit swift behavioral reactions to humidity changes [48]. While our SBF-SEM analysis reveals humidity-dependent structural differences, these represent static snapshots of what is likely a highly dynamic process occurring on a millisecond timescale [41]. The ability to capture these rapid structural changes presents a technical challenge, as current methodologies require tissue fixation that precludes temporal analysis.

The requirement for rapid humidity detection has likely shaped the evolution of hygrosensilla across insect species, resulting in diverse structural adaptations that balance sensitivity with response time [14,15]. In cockroaches, "sensilla capitulum" are shielded by bristles and a cuticular wall, reducing their exposure to air pressure changes [49,50]. Conversely, stick insects have a more exposed "peg-in-pit" structure surrounded by a protective collar, allowing direct environmental interaction [51]. These adaptations suggest species-specific optimization between exposure and protection. In *D. melanogaster*, the location of hygrosensilla within the sacculus presents an intriguing paradox. This invagination on the posterior antenna creates a protected environment but may also limit the diffusion of air and potentially slow sensory responses. We propose that passive diffusion of air alone may be insufficient to support the observed rapid behavioral responses to humidity changes. We suggest that active airflow within the sacculus, potentially facilitated by antennal movements, could enhance humidity detection. Although speculative, this concept parallels observations in species such as cockroaches, where antennal fanning enhances olfactory sensitivity, and may warrant investigation in *D. melanogaster* [52,53]. This model would reconcile the apparently contradictory requirements for protection and rapid response.

The chamber-specific structural adaptations we observe may further enhance this active sensing system. The sharper tapering of sensilla under different humidity conditions could create localized regions of enhanced mechanical sensitivity, optimizing detection despite the constraints of the sacculus environment. This specialized architecture, combined with the proposed active airflow mechanism, could enable the rapid and precise humidity detection necessary for adaptive behavior. Future studies combining structural analysis with real-time measurements of sensilla dynamics could provide valuable insights into how these spatial and temporal aspects of humidity sensing are integrated.

### Concluding remarks

Our study demonstrates how combining SBF-SEM and deep learning-based segmentation reveals fundamental aspects of sensory biology, providing the first three-dimensional analysis of humidity-dependent structural changes if insect hygrosensilla. While identifying humidity-induced structural adaptations in *D. melanogaster* hygrosensilla, several important questions remain to be addressed, including the temporal dynamics of hygrosensilla structural changes, the molecular mechanisms underlying the observed structural plasticity, and how different evolutionary pressures have shaped hygrosensilla architecture across species. The methodological pipeline we have developed, combining precise environmental control, rapid preservation, high-resolution imaging, and machine learning-based analysis, provides a powerful approach that can be extended to study structural changes in other sensory organs, establishing a technical framework for future studies of sensory structure-function relationships.

## Acknowledgments

The authors thank Klaus Qvortrup, Tillmann Pape, Zhila Nikrozi and Cristiano Di Benedetto at the Core Facility for Integrated Microscopy, Faculty of Health and Medical Sciences, University of Copenhagen, for technical assistance with the sample preparation and imaging.

## Author Contribution

AE and GG designed the study. AE and GG performed all experiments. GG performed all data analysis. AE and GG wrote the manuscript.

## Data availability

Data and scripts used for analysis can be found on https://github.com/enjinlab/SacculusUltrastructure.

